# Identification of a novel population of peripheral sensory neuron that regulates blood pressure

**DOI:** 10.1101/2020.01.17.909960

**Authors:** Chiara Morelli, Laura Castaldi, Sam J. Brown, Lina L. Streich, Alexander Websdale, Francisco J. Taberner, Blanka Cerreti, Alessandro Barenghi, Kevin M. Blum, Julie Sawitzke, Tessa Frank, Laura Steffens, Balint Doleschall, Joana Serrao, Stefan G. Lechner, Robert Prevedel, Paul A. Heppenstall

## Abstract

The vasculature is innervated by a network of peripheral afferents that sense and regulate blood flow. Here, we describe a system of non-peptidergic sensory neurons with cell bodies in the spinal ganglia that regulate vascular tone in the distal arteries. We identify a population of mechanosensitive neurons marked by TrkC and Tyrosine hydroxylase in the dorsal root ganglia that project to blood vessels. Local stimulation of these neurons decreases vessel diameter and blood flow, while systemic activation increases systolic blood pressure and heart rate variability via the sympathetic nervous system. Chemogenetic inactivation or ablation of the neurons provokes variability in local blood flow leading to a reduction in systolic blood pressure, increased heart rate variability and ultimately lethality within 48 hours. Thus, TrkC/Tyrosine hydroxylase positive sensory neurons form part of a sensory feedback mechanism that maintains cardiovascular homeostasis through the autonomic nervous system.

## Introduction

Regulation of blood pressure through control of blood flow and cardiac output is essential for tissue and organ homeostasis. This is achieved through local mechanisms intrinsic to blood vessels such as endothelial cell derived signaling, and via neuronal mechanisms that regulate cardiovascular function at a global level to coordinate vascular resistance and modulate cardiac output (Thomas, 2011; Westcott and Segal, 2013). Indeed, major blood vessels are surrounded by a dense plexus of nerves made up of post-ganglionic sympathetic efferent axons and sensory axons from the spinal ganglia. These exert broadly opposing effects on vascular resistance such that activation of sympathetic neurons causes vasoconstriction, while activity in sensory nerves produces vasodilation.

Two types of perivascular sensory neuron have been previously described: peptidergic vascular afferents, which originate in the dorsal root ganglia (DRG), and mechanosensitive baroreceptors of the nodose and petrosal ganglia (Coleridge and Coleridge, 1980; Westcott and Segal, 2013). Peptidergic sensory neurons can be identified by their expression of neuropeptides, such as Substance P (SP) and Calcitonin Gene Related Peptide (CGRP), and the ion channel TRPV1 (Gibbins et al., 1985; Vass et al., 2004). They are activated by mechanical, chemical, and thermal stimuli (Adelson et al., 1997; Bessou and Perl, 1966; Haupt et al., 1983), and project to vascular beds throughout the body (Burnstock and Ralevic, 1994). As well as having an afferent sensory function, it has long been appreciated that these neurons also perform an effector role through local axon reflexes (Bayliss, 1901; Bruce, 1913). Thus their activation by noxious stimuli in tissue can provoke a potent vasodilator effect through the release of CGRP and purines (Burnstock, 2007; Kawasaki et al., 1988), as well as plasma extravasation through SP (Lembeck and Holzer, 1979).

Baroreceptors are stretch-sensitive sensory neurons that project from the cervical ganglia to the aorta and carotid sinuses (Kirchheim, 1976). Their endings are embedded in artery walls and they continuously monitor arterial blood pressure through the mechanosensitive ion channels Piezo1 and Piezo2 (Zeng et al., 2018). They are exquisitely sensitive to distension of the vessel wall such that their firing rate fluctuates as arterial pressure changes through the cardiac cycle. They function to minimize short-term fluctuations in blood pressure; thus, increases in baroreceptor activity provoke arteriolar vasodilation through inhibition of the sympathetic nervous system, and reduction in cardiac output via decreased sympathetic and increased parasympathetic input. Conversely, decreased baroreceptor activity produces vasoconstriction and increased cardiac output (Wehrwein and Joyner, 2013). Both clinical and experimental studies have demonstrated that a reduction in baroreceptor function results in a marked increase in lability of blood pressure and heart rate (Heusser et al., 2005; Ito and Scher, 1981; Robertson et al., 1993; Rodrigues et al., 2011; Sved et al., 1997).

Beyond peptidergic vascular afferents and baroreceptors, further classes of perivascular sensory neurons are likely to exist, especially those involved in sensing vessel wall stretch. However, their identification has been hampered by their inaccessibility to classical neurophysiological methods, and by a paucity of molecular markers by which to distinguish them. Here, we identify a population of non-peptidergic DRG neurons that coexpress TrkC and Tyrosine Hydroxylase and project to distal arteries. We demonstrate that experimental ablation of these neurons leads to dysregulation of blood flow and heart rate, decreased blood pressure, and ultimately death of mice within 48 hours. Moreover, their optogenetic or chemogenetic activation provokes vasoconstriction, marked increases in blood pressure and heart rate variability that are dependent upon activation of the sympathetic nervous system, and pain behavior. Thus, we describe a new population of vascular sensory afferents that together with peptidergic afferents and baroreceptors coordinate vascular resistance and cardiac output.

## Results

To characterize new subpopulations of peripheral neuron we generated an inducible TrkC^CreERT2^ mouse line and crossed it with a Rosa26^ChR2-YFP^ reporter line to examine the colocalization of TrkC with established cellular markers in adult peripheral ganglia. We first investigated the DRG, where TrkC has been previously shown to colocalize with parvalbumin (Pvalb, a marker of proprioceptors (Copray et al., 1994)), and Ret (as a marker of Aβ Field mechanoreceptors (Bai et al., 2015)). 31% of all DRG neurons were positive for TrkC^CreERT2^-driven YFP. Of these, 31% were labelled with Pvalb and NF200 (Figures 1A and 1F) and 38% co-expressed Ret and NF200 (Figures 1B and 1F), while 31% of TrkC^+^ neurons expressed neither of these markers, nor NF200 (Figure 1F). To identify this third population, we stained DRG sections for markers of non-peptidergic nociceptors (IB4), peptidergic nociceptors (CGRP), and C fiber low-threshold mechanoreceptors (C-LTMRs, Th (Li et al., 2011)). We observed minimal overlap between TrkC^CreERT2^-driven YFP and IB4 (Figures 1C and 1F) and CGRP (Figures 1D and 1F), but found that most TrkC^+^/NF200^-^ neurons expressed Th (Figures 1E and 1F). TrkC^+^/Th^+^ neurons were significantly smaller than TrkC^+^/Th^-^ neurons (Figure 1G); they were also more prevalent in lumbar DRG than thoracic or cervical DRG (Figure 1H; Figures S1A-S1C). We further investigated whether TrkC^+^/Th^+^ neurons were evident in other peripheral ganglia. We observed robust Th labelling in the nodose-petrosal-jugular ganglion complex, but no TrkC^CreERT^-mediated recombination (Figure 1I; Figures S1D and S1E). Similarly, sympathetic neurons in the paravertebral ganglia were positive for Th, but negative for TrkC (Figure 1J; Figures S1F and S1G).

**Figure 1.**
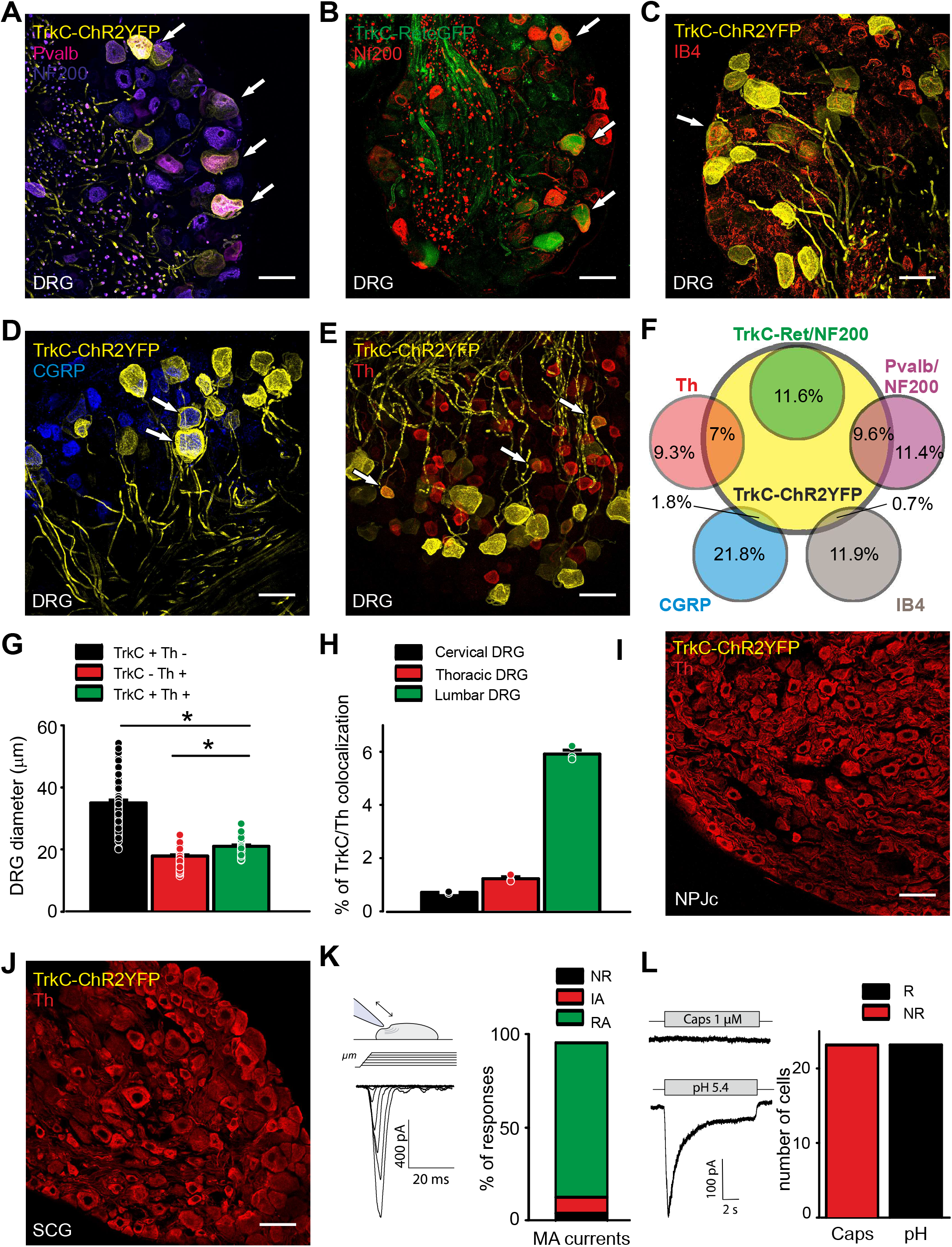
TrkC expression in peripheral ganglia. Immunofluorescent staining of DRG sections from TrkC^CreERT2^::Rosa26^ChR2-YFP^ showed that 9.6% of DRG neurons co-express markers of proprioceptors (PValb/NF200) (A and F) and 11.6% of SA field mechanoreceptors (Ret/NF200) (B and F). Very few TrkC^+^ neurons overlap with markers of non-peptidergic nociceptors (IB4) (C and F), or peptidergic nociceptors (CGRP) (D and F). 7% of DRG neurons are also positive for Th, a marker of C fiber low-threshold mechanoreceptors (C-LTMRs) (E and F). (F) Quantification of TrkC co-expression with the other markers in DRG showing the percent of total neurons. (G)TrkC^+^/Th^+^ DRG neurons are significantly larger than TrkC^-^/Th^+^ neurons and smaller than TrkC^+^/Th^-^ neurons (298 neurons from 3 mice, *p<0,001). (H) Quantification of the number of DRG neurons coexpressing TrkC and Th at different spinal segmental levels, expressed as percentage of the total number of DRG neurons (n=3). (I and J) TrkC is not expressed in the nodose-petrosal-jugular ganglion complex (I), nor in the superior cervical ganglion (J). (K) *Left*, sketch depicting the mechanical stimulation protocol and an example trace of a mechanically activated current (MA current) in a small TrkC^+^ neuron. *Right*, distribution of the different types of MA currents in the small TrkC^+^ neurons (24 neurons from 3 mice): rapidly adapting (RA), intermediate adapting (IA), non-responding (NR). (L) *Left*, Example responses to capsaicin and pH 5.4 in a small TrkC^+^ neuron. *Right*, Number of neurons responding (R) and non-responding (NR) to capsaicin or pH in small TrkC^+^ neurons. Arrows indicate co-expression of TrkC^+^ neurons with the other markers. Scale bars, 50μm.

We next explored the response properties of isolated small TrkC^+^ DRG neurons using wholecell patch clamp. All neurons were responsive to mechanical stimulation and displayed a rapidly inactivating current indicative of Piezo2 activation (Figure 1K). Neurons were also activated by low pH, but were not sensitive to the TRPV1 agonist capsaicin (Figure 1L; Figures S1H-S1J). We further analyzed published single cell RNAseq data of DRG neurons (Zeisel et al., 2018) and consistent with our findings, distinguished a population of TrkC^+^/Th^+^ neurons (46% of all TrkC neurons, 65% of all Th neurons) that expressed high levels of Piezo2 and the acid-sensing ion channel ASIC2, but were negative for Parvalbumin, TRPV1, and CGRP (Figure S1K).

Given that Th^+^ C-LTMRs project to the skin and form longitudinal lanceolate endings around hair follicles (Li et al., 2011), we examined whole mount preparations of skin for TrkC^CreERT2^-driven YFP fluorescence. While we observed clusters of TrkC^+^ Aβ Field mechanoreceptors (Figure 2A), we were unable to detect any longitudinal lanceolate endings indicative of C-LTMRs. We did however discern large spiral structures that extended for several millimeters along blood vessels (Figure 2A). Further characterization of these TrkC^+^ structures revealed that they surrounded arterioles throughout the body (Figures 2B-2D). The spirals were also negative for desmin (Figures 2E-2G), a marker of pericytes, suggesting that they are in fact vascular smooth muscle cells (vSMCs).

**Figure 2.**
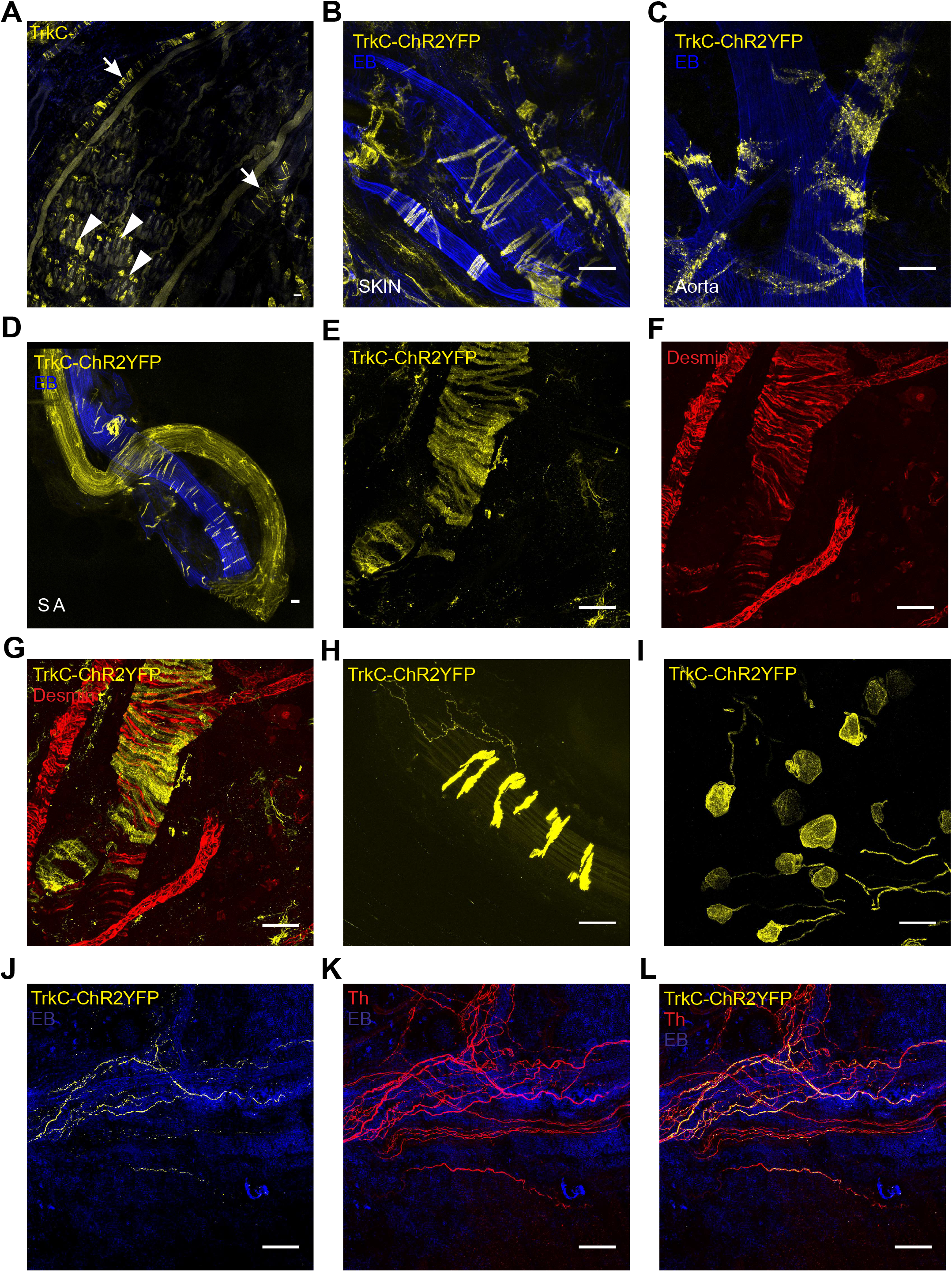
TrkC mediated recombination in peripheral tissue. (A) A whole mount skin preparation from a TrkC^CreERT2^::Rosa26^ChR2-YFP^ mouse shows TrkC^+^ Aβ Field mechanoreceptors (arrowheads) and TrkC^+^ structures surrounding blood vessels (arrows).(B-D) TrkC is expressed in vSMCs throughout the body: skin (B), aorta (C), saphenous artery (D). Mice were injected with Evans Blue (EB) i.v. to visualize blood vessels. (E-G) Immunohistochemical analysis with anti-desmin antibodies in TrkC^CreERT2^::Rosa^26ChR2-YFP^ mice reveals that TrkC does not mark pericytes but vSMCs. (H) TrkC^+^ vSMCs are innervated by TrkC^+^ neurons. (I-L) Intrathecal injection of 4-hydroxytamoxifen in TrkC^CreERT2^::Rosa26^ChR2-YFP^ mice leads to robust expression of YFP in DRG (I) and in neurons innervating blood vessels (J), but not in vSMCs (J). (K and L) TrkC^+^ neurons innervating the vessels co-express the marker Th. Scale bars, 50μm.

Intriguingly, upon examination of vessels at higher magnification, it became evident that they were innervated by TrkC^+^ neurons (Figure 2H). To visualize these afferents in the absence of contaminating YFP fluorescence from TrkC^+^ vSMCs, we injected the active tamoxifen metabolite 4-hydroxytamoxifen intrathecally in TrkC^CreERT2^ mice to induce Cre expression only in DRG (Figures S3A-S3L). We observed robust recombination in DRG (Figure 2I; Figures S2A-S2F) and clearly visible innervation of vessels (Figure 2J and 2L;Figure S2G-S2L). Costaining with Th revealed that all TrkC^+^ vessel innervating afferents were Th^+^ and that they were interweaved with presumed sympathetic neurons, forming a dense plexus around vessels (Figure 2J-2L).

To determine the function of TrkC^+^/Th^+^ neurons, we first took a loss of function approach using TrkC^CreERT2^::Avil^iDTR^ mice to delete TrkC^+^ neurons solely in adult sensory ganglia. Unexpectedly, we observed 100% lethality in these mice within 48 hours after injection of diphtheria toxin (DTX) (Figure 3A). To assess the extent of ablation, in particular its specificity to sensory neurons, we generated triple transgenic TrkC^CreERT2^::Avil^iDTR^Rosa26^ChR2-YFP^ to visualize the loss of TrkC^+^ cells. Following DTX injection, we detected a complete absence of TrkC^+^ neurons in the DRG (Figure 3B) and innervating blood vessels (Figure 3C). Importantly, however, Th^+^ fibers were still evident around vessels (red signal, Figure 3C), likely representing sympathetic neurons, as were TrkC^+^ vSMCs (yellow signal, Figure 3C).

**Figure 3.**
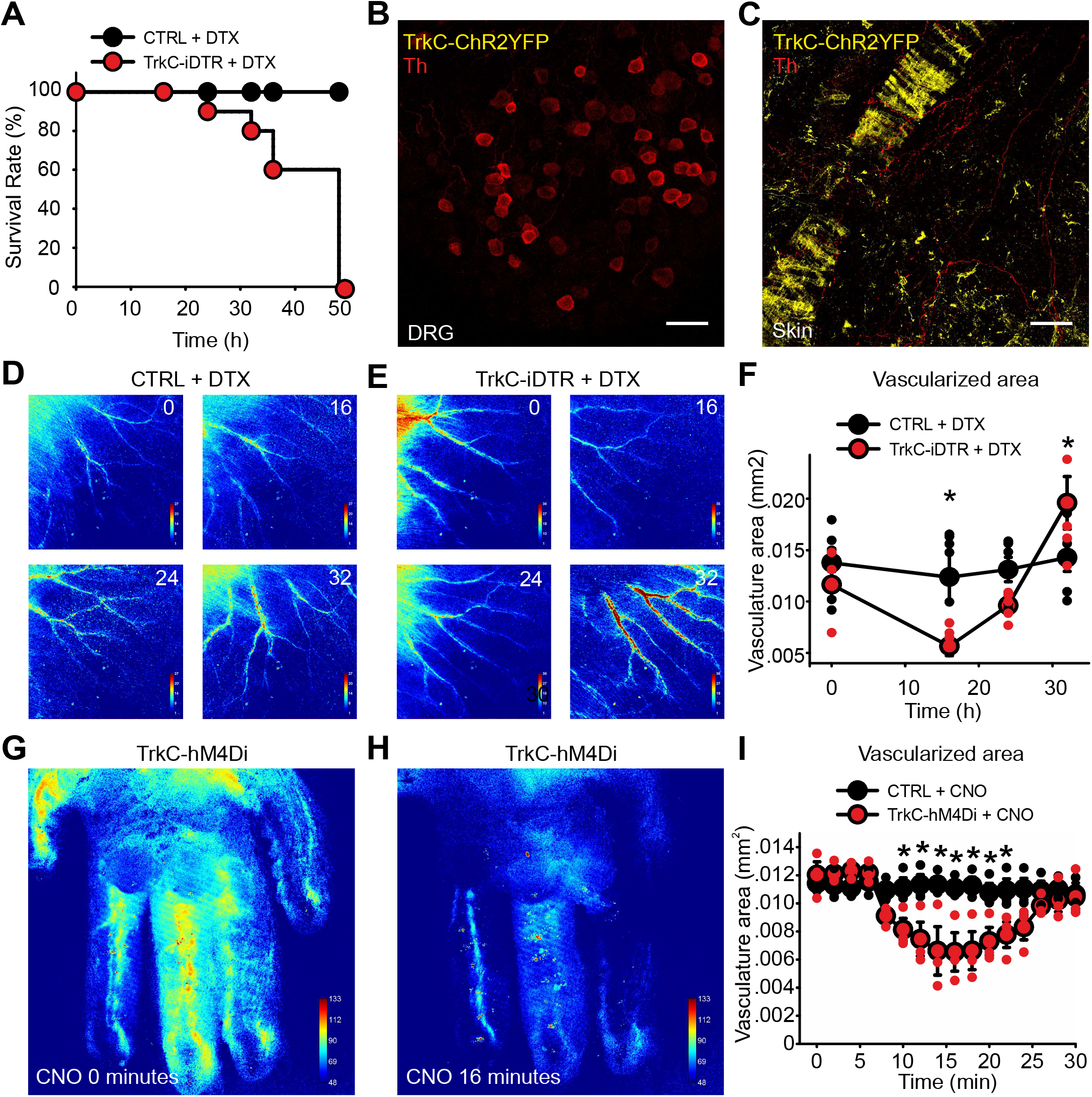
Loss of function in TrkC^+^ neurons. (A) Survival rate of TrkC^CreERT2^Avil^iDTR^ (red symbols, n=10) and control mice (black symbols, n=10) upon intraperitoneal administration of DTX. (B and C) Ablation leads to a complete absence of TrkC^+^ neurons in DRG and of TrkC^+^/Th^+^ fibers innervating blood vessels. Immunofluorescence with Th antibodies on DRG sections (B) and whole mount skin (C) from TrkC^CreERT2^::Avil^iDTR^::Rosa26^ChR2-YFP^ mice after DTX injection. Scale bars, 50μm. (D and E) Representative LSCI images of the ear of a control (D) and a TrkC^CreERT2^::Avil^iDTR^ mouse (E) before DTX injection (0) and at 16, 24, 32 hours after treatment. (F) Measurements of the vascularized area showed blood flow alteration upon ablation of TrkC^+^ neurons (*p<0,05). (G and H) Representative LSCI images showing blood vessels in the hind paw of a TrkC^CreERT2^::AAV-PHP.S^hM4Di^ mouse before treatment (G) and 16 min after CNO injection (H). (I) Measurements of the vascularized area showed blood flow alteration upon inhibition of TrkC^+^ neurons with CNO (*p<0,05, n=3).

We next investigated the impact of TrkC^+^ neuron loss on local blood flow using Laser Speckle Contrast Imaging (LSCI) in anesthetized mice. Sixteen hours after DTX injection, there was a significant reduction in peripheral blood flow in the skin of mice (Figures 3D and 3E) to the extent that smaller arteries were no longer detectable (reduction in vascularized area, Figure 3F) and relative perfusion in larger vessels was also reduced. Over the next 16 hours, this then rebounded such that shortly before death, mice displayed substantially higher blood flow (Figures 3E and 3F). To further explore the consequences of short-term inactivation of these neurons, we performed LCSI on TrkC^CreERT2^ mice infected with an AAV-PHP.S virus (Chan et al., 2017) carrying the inhibitory Gi coupled DREADD receptor hM4Di under a neuronal specific promoter (Krashes et al., 2011). Upon local injection of the DREADD agonist clozapine N-oxide (CNO) into the paw, we again observed a significant reduction in vascularized area that maximized within 15 minutes and recovered to baseline values by 30 minutes (Figures 3G-3i).

To investigate factors that may be contributing to lethality upon ablation of TrkC^+^ neurons, we monitored systolic blood pressure and heart rate after injection of DTX. At 16 hours post-DTX injection, we observed a large decrease in blood pressure (107±6 mm Hg compared to 133±4 mm Hg before injection) that further decreased at 24 hour and 32 hour time points (24 hours: 100±7 mm Hg, 32 hours: 96±6 mm Hg) (Figure 4A). Heart rate measurements showed a complex progression over the 48 hours. Average heart rate was not different between treated mice and controls at any time point (Figure 4B). However, increases in heart rate variability (HRV) were clearly apparent at 24 (Figure 4C) and 32 hours after DTX injection (Figure 4D).

This was evidenced by a significant increase in long-term variability over 30 minutes (Figure 4E; Figures S3A and S3B), but not in beat-to-beat, short-term variations in the signal (Figure 4F), suggesting autonomic dysregulation (Electrophysiology Task Force of the European Society of Cardiology the North American Society of, 1996).

**Figure 4.**
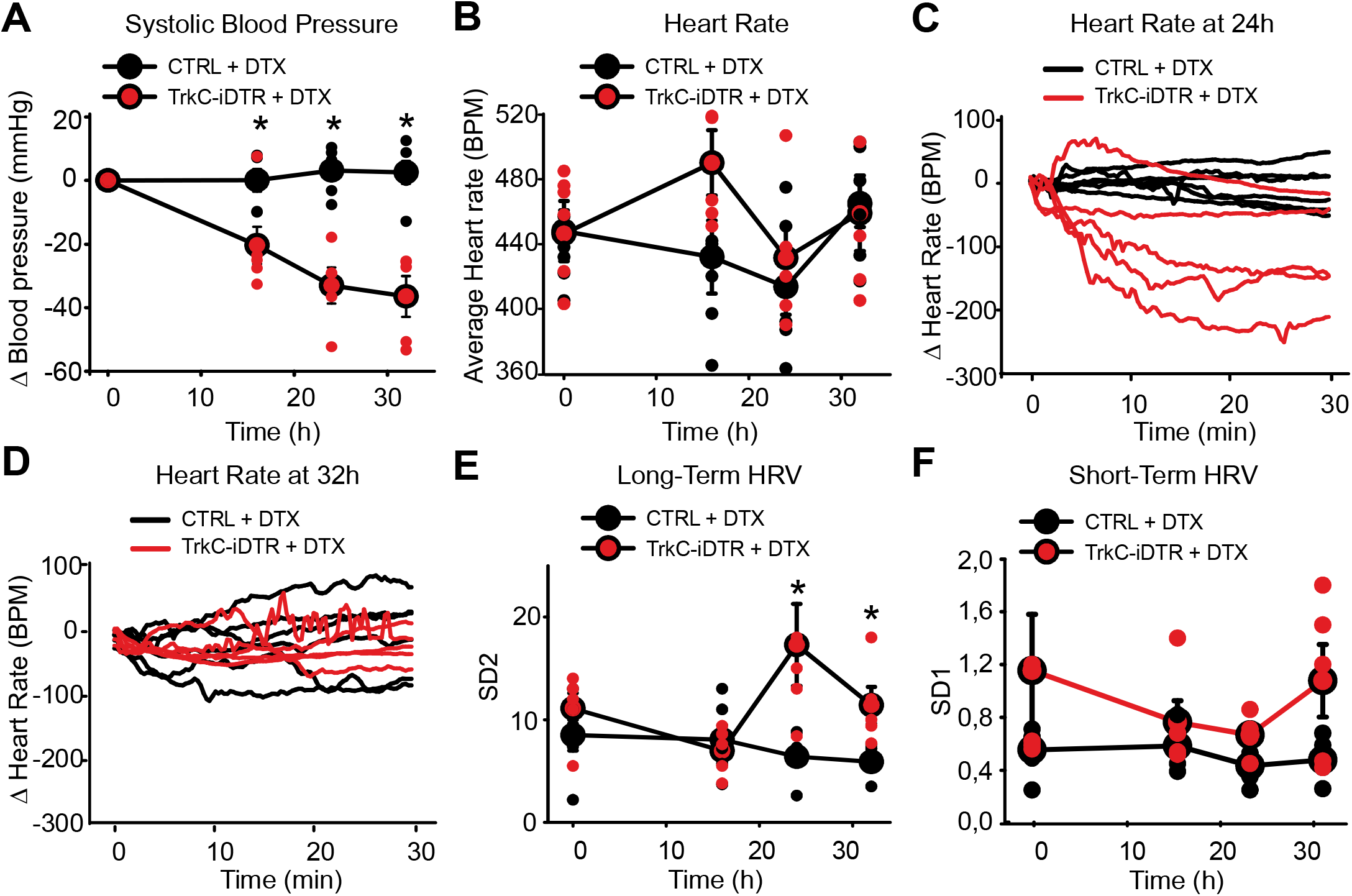
Ablation of TrkC^+^ neurons leads to blood pressure and heart rate alterations. (A) Upon systemic injection of DTX, TrkC^CreERT2^::Avil^iDTR^ mice display a decrease in blood pressure (red symbols, n=5), while control mice (black symbols, n=6) are not affected (*p<0,001). (B) Average heart rate in TrkC^CreERT2^::Avil^iDTR^ mice (red symbols, n=5) after DTX injections at different time points over 30 minutes compared with controls (black symbols, n=6). (C and D) Fluctuations in heart rate for individual mice over 30 minutes at 24 (c) and 32 (d) hours after DTX injection. (E) Long-term heart rate variability derived from the length of the major (SD2) axis of Poincaré plots at different time points after injection of DTX. Significant differences in variability are evident in TrkC^CreERT2^::Avil^iDTR^ mice compared to controls at 24h and 32h post-injection (*p<0,05). (F) Short-term variability derived from the length of the minor (SD1) axis of Poincaré plots at different time points after injection of DTX in TrkC^CreERT2^::Avil^iDTR^ and control mice.

To investigate the mechanism by which TrkC^+^/Th^+^ sensory neurons regulate the cardiovascular system, we took a gain-of-function approach to activate TrkC^+^ neurons, and monitored blood flow, vessel diameter, blood pressure, and heart rate upon stimulation. We utilized an Avil^hM3Dq^ mouse line that allows for Cre-dependent expression of the excitatory Gq coupled DREADD receptor only in peripheral neurons (Dhandapani et al., 2018b). To monitor blood flow at the macroscopic tissue level, we injected CNO into the paw of anesthetized TrkC^CreERT2^::Avil^hM3Dq^ mice and imaged the vasculature for 30 minutes using LSCI (Figure 5A). Ten minutes post-injection, a significant reduction in vascularized area and perfusion rate was evident that progressively decreased such that after 20 minutes the vasculature became essentially undetectable (Figures 5B and 5C). To examine this further at the microscopic single-vessel level, we performed *in-vivo* three-photon (3P) microscopy in conjunction with optogenetic activation of TrkC^+^ neurons in the ear of anesthetized TrkC^CreERT2^::Rosa26^ChR2-YFP^ mice injected intrathecally with 4-hydroxytamoxifen. Intriguingly, upon optical stimulation, we observed a significant decrease in vessel cross-sectional area in TrkC^CreERT2^::Rosa26^ChR2-YFP^ mice that was not apparent in control animals (Figures 5D and 5E; Video S1). To confirm that this was associated with a reduction in blood flow as seen in LCSI experiments, red blood cell velocity was calculated from spatially windowed 3P line-scan data. As shown in Figures 5F and 5G, Video S2 and S3, optical stimulation provoked a dramatic and persistent decrease in blood flow, such that individual red blood cells became clearly visible at the imaging frame rate.

**Figure 5.**
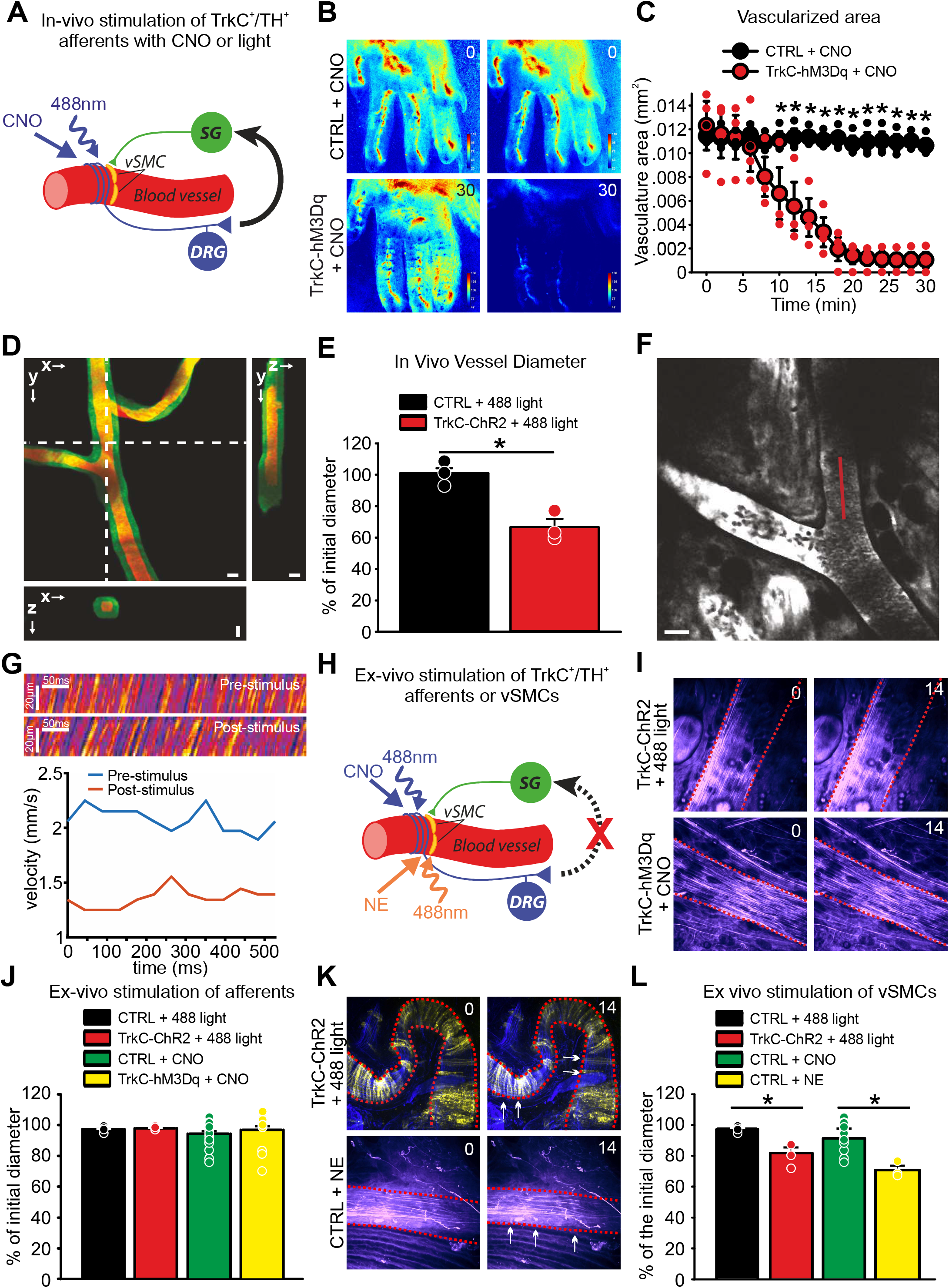
Activation of TrkC^+^ neurons. Schematic showing circuit activated by CNO or 488 nm light in vivo. (B) Representative LSCI images showing blood vessels in the hind paw of control mice (upper panels) or TrkCCreERT2::AvilhM3Dq mice (lower panels) treated locally with CNO imaged before and 30 min after the treatment. (C) Measurements of the vascularized area of LSCI images showing blood flow reduction upon local activation of TrkC^+^ neurons with CNO (red symbols, *p<0,001, n=3). (D) 3-photon volumetric image of blood vessels in the mouse ear labeled with dextran-fluorescein, Maximum intensity projection along lateral imaging plane (x/y) and orthogonal projection of 3D volume along lines indicated in (x/y) image by the white dotted line. Image of green labeled artery was acquired before the optogenetic stimulus and red after the stimulus. Scale bar 25 μm. (E) Quantification of blood vessel diameter after optical stimulation (*p<0.005). (F) 3-photon image of blood vessels in the mouse ear labeled with dextran-fluorescein. Red line indicates the laser scan path used to measure and calculate red blood cell velocity. Line-scans were acquired at 666Hz. (G) Upper: line scans generated from the path indicated by the red line in (F) can be stacked as a space-time (x/t) plot, where apparent angle is proportional to flow velocity. The images show ~500ms of data collection before and after optogenetic stimulation, respectively. Lower: red blood cell velocity before and after stimulation. Velocity was calculated based on Radon-analysis of spatially windowed space-time plot data (window size 50ms). (H) Schematic showing stimulation of TrkC^+^/Th^+^ afferents or vSMCs ex vivo. (I) Representative images of blood vessels from an ex vivo skin-nerve preparation prior to and 14 minutes after optogenetic activation of TrkC^+^ neurons (top panels) or application of CNO (lower panels). (J) Quantification of blood vessel diameter after ex vivo stimulation of TrkC^+^ afferents with 488 nm light or CNO application. (K) Representative images of blood vessels prior to and 14 minutes after optogenetic activation of vSMCs (top panels) or application of norepinephrine (lower panels). Dotted lines indicate blood vessel perimeter, arrows indicate shrinkage. (L) Quantification of blood vessel diameter after ex vivo stimulation of vSMCs with 488 nm light or norepinephrine application (*p<0.01).

To investigate whether TrkC^+^/Th^+^ neurons exert their effects on blood vessels locally, we next performed confocal live imaging in an isolated ex vivo preparation of the hind limb skin where spinal reflex arcs were disrupted (Figure 5H). We never detected changes in vessel size upon activation of TrkC^+^ neurons using either chemogenetic or optogenetic stimulation (Figures 5I and 5J; Video S4 and S5). However, direct activation of vSMCs expressing ChR2 using optogenetics, or via application of norepinephrine, provoked robust constriction of vessels indicating that the preparation was viable (Figures 5K and 5L; Video S6 and S7). Thus, TrkC^+^/Th^+^ neurons are not likely to exert an efferent function via axon reflex, nor do they appear to act through local connections with sympathetic neurons.

We further assessed the influence of TrkC^+^ neuronal activation on systolic blood pressure and heart rate (Figure 6A). We used the DREADD agonist 21 (C21) (Chen et al., 2015) to avoid off-target systemic effects associated with metabolism of CNO to clozapine (Gomez et al., 2017). C21 was injected intraperitoneally in TrkC^CreERT2^::Avil^hM3Dq^ mice, and within 10 minutes, we observed a substantial increase in blood pressure from 117±3 mm Hg to 154±2 mm Hg that persisted for 30 minutes (Figure 6B). Average heart rate also increased significantly over the same time course (Figure 6C) and heart rate variability became evident (Figure 6D). This was accompanied by an increase in long-, but not short-term, measures of HRV (Figures 6E and 6F; Figures S4A and S4B). We next treated mice with the β-adrenergic receptor–blocker propranolol to inhibit sympathetic input (Figure 6A). In the presence of propranolol, C21 did not provoke any changes in blood pressure (Figure 6B), heart rate (Figures 6C and 6D), or variability (Figures 6E and 6F; Figures S4C and S4D), indicating that TrkC^+^/Th^+^ neurons signal via the sympathetic nervous system to regulate cardiovascular output.

**Figure 6.**
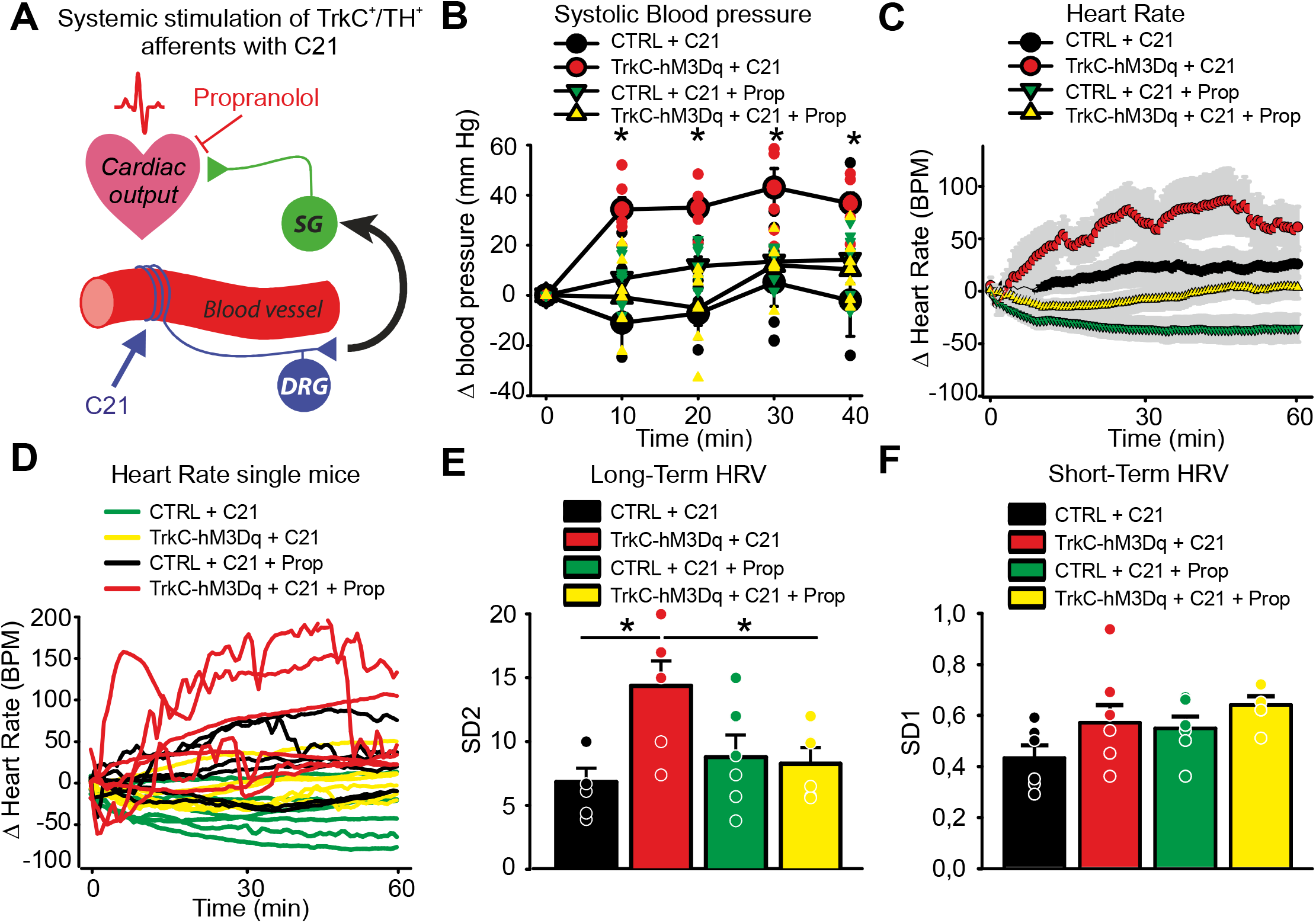
Systemic chemogenetic activation of TrkC^+^ neurons leads to blood pressure and heart rate alterations. (A) Schematic showing circuit activated by C21 in vivo. (B) Systemic injection of C21 in TrkC^CreERT2^::Avil^hM3Dq^ mice leads to elevated blood pressure (red symbols, n=6) compared to control mice (black symbols, n=6), *p<0,01. This is reverted by co-administration of propranolol (yellow symbols, n=6). (C) TrkC^CreERT2^::Avil^hM3Dq^ mice (red symbols, n=6) treated systemically with C21 display increased average heart rate compared to mice co-treated with propranolol (yellow symbols, n=6) or control mice (black and green symbols). (D) Fluctuations in heart rate in individual mice are apparent in TrkC^CreERT2^::Avil^hM3Dq^ mice treated with C21 (red lines), but not in mice co-treated with propranolol (yellow lines) or control animals (black and green lines)., (E) Long-term heart rate variability derived from the length of the major (SD2) axis of Poincaré plots in TrkC^CreERT2^::Avil^hM3Dq^ mice treated with C21, C21 plus propranolol, or in control mice (*p<0,05). (F) Short-term heart rate variability derived from the length of the minor (SD1) axis of Poincaré plots in TrkC^CreERT2^::Avil^hM3Dq^ mice treated with C21, C21 plus propranolol, or in control mice.

Finally, we investigated the consequences of local activation of TrkC^+^ neurons and subsequent reductions of peripheral blood flow, on sensory behavior. CNO was injected subcutaneously in the paw and responses to mechanical and thermal stimulation of the skin were monitored (Figure 7A). We observed a striking reduction in mechanical thresholds after CNO injection (Figure 7B), such that 10 minutes post-injection, mice became hypersensitive to the lightest of punctate stimuli (Figure 7C). Responses to dynamic brushing stimuli and thermal stimuli were not altered by CNO (Figures 7D-7F).

**Figure 7.**
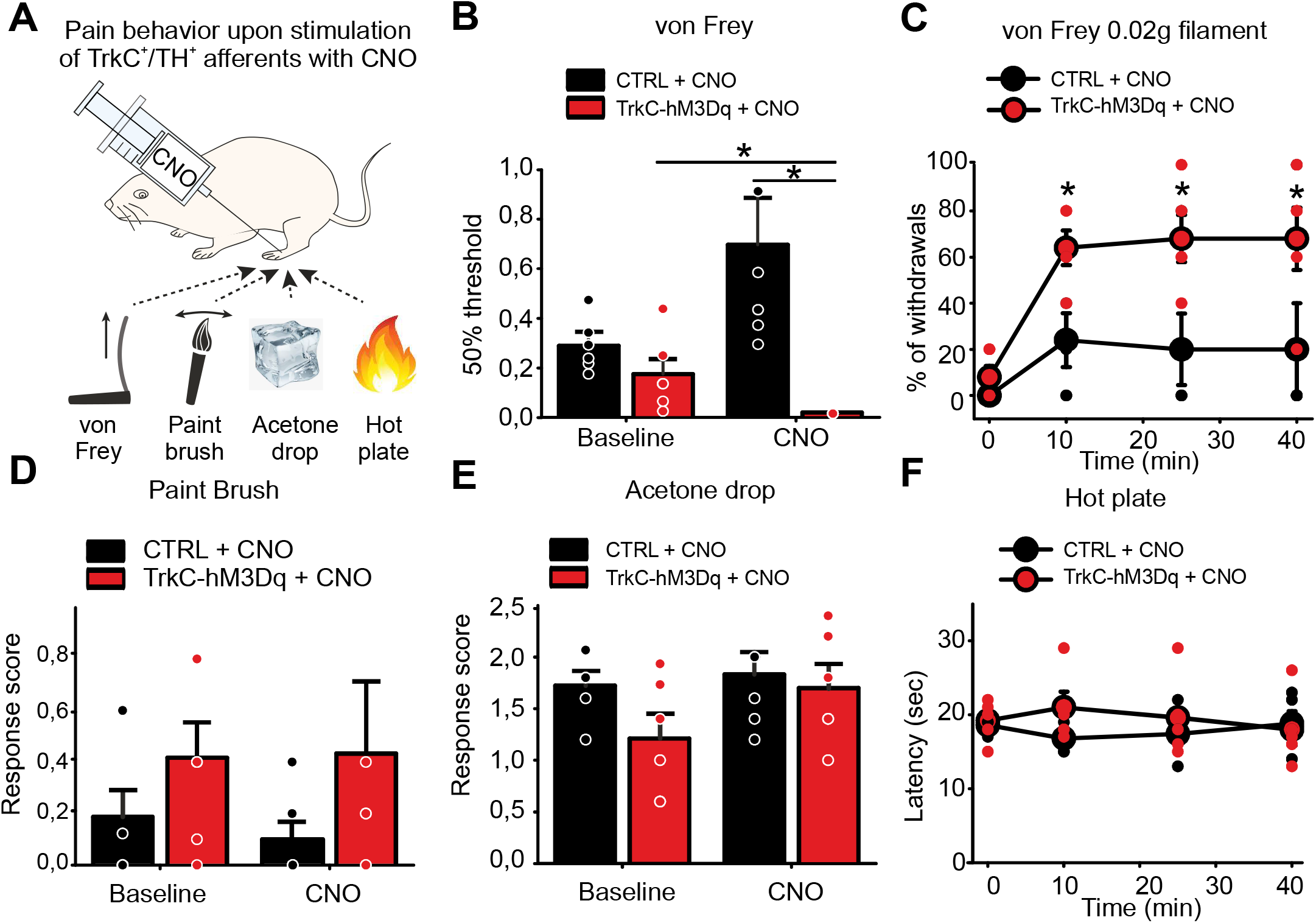
Behavioral responses to mechanical and thermal stimulation upon chemogenetic activation of TrkC^+^ sensory neurons. (A) Schematic of behavioral tests performed after local injection of CNO. (B and C) TrkC^CreERT2^::Avil^hM3Dq^ developed hypersensitivity to punctate mechanical stimuli (n=6, *p<0.05) (B) that started 10 min after CNO local treatment and persisted for more than 40 minutes (C) (0.02g filament, n=5, *p<0.05). (D) Local intraplantar injection of CNO in TrkC^CreERT2^::Avil^hM3Dq^ mice does not provoke mechanical hypersensitivity to dynamic mechanical stimuli (paintbrush, n=6). (E and F) Thermal sensitivity as assayed by the acetone drop test (n=6) (E) and hot plate test (n=5) (F) was not different between TrkC^CreERT2^::Avil^hM3Dq^ (red symbols) and control mice (black symbols) treated with CNO.

## Discussion

Here, we describe a population of peripheral neurons marked by TrkC and Th that have cell bodies in the spinal ganglia and project to distal blood vessels. We demonstrate that silencing or ablation of these neurons leads to rapid reductions in blood flow to the skin, and that over the course of 48 hours this is accompanied by a catastrophic drop in blood pressure and increased heart rate lability that ultimately results in the death of mice. Moreover, we show that activation of these neurons provokes vasoconstriction, and increased blood pressure and heart rate variability. This is dependent upon a spinal or supra-spinal reflex arc that requires β-adrenergic signaling, implicating a role for the sympathetic nervous system. Intriguingly, activation of the neurons also provokes a profound and rapid hypersensitivity to punctate mechanical stimulation of the skin.

Our data indicate that TrkC^+^/Th^+^ neurons are a discrete population of DRG neuron that are molecularly, anatomically, and functionally distinct from previously described perivascular sensory neurons. In contrast to peptidergic neurons, they do not express CGRP, are not activated by capsaicin, and have no apparent efferent function through axon reflexes (Bayliss, 1901; Bruce, 1913; Gibbins et al., 1985; Vass et al., 2004). Upon stimulation they also provoke vasoconstriction rather than vasodilation (Burnstock, 2007; Kawasaki et al., 1988). Thus, functionally they more closely resemble baroreceptors, and indeed, they appear to contribute to a baroreceptor reflex-like arc to control arterial pressure. However, baroreceptors have their cell bodies in the cervical ganglia and project to the aortic arch and carotid artery (Kirchheim, 1976), while TrkC^+^/Th^+^ neurons are located in the spinal ganglia and project to distal arteries. Moreover, baroreceptor stimulation provides negative feedback to cardiovascular output (Coleridge and Coleridge, 1980), while activation of TrkC^+^/Th^+^ neurons has opposing effects and raises blood pressure and heart rate. Importantly, these differences are also apparent in ablation studies. Thus, depletion of peptidergic vascular afferent nerves by repeated exposure to capsaicin does not disrupt baroreceptor reflexes (Furness et al., 1982), while surgical denervation of baroreceptors results in a marked increase in blood pressure variability often accompanied by an increase in arterial blood pressure (Heusser et al., 2005; Ito and Scher, 1981; Robertson et al., 1993; Rodrigues et al., 2011; Stocker et al., 2019; Sved et al., 1997). In contrast, our data indicates that DTX mediated ablation of TrkC^+^/Th^+^ neurons leads to a significant decrease in systolic blood pressure, accompanied by death of mice within 48 hours. We thus speculate that TrkC^+^/Th^+^ neurons are essential for integrating information on vascular status from distal tissues and providing positive feedback to the cardiovascular system. Investigating how these neurons signal and respond under different physiological conditions, and determining whether they exist in other species beyond mice are important questions for future studies.

In addition to its expression in TrkC^+^/Th^+^ neurons, Th is also a marker of sympathetic neurons where it functions as the enzyme that catalyzes the biosynthesis of catecholamines, such as norepinephrine. This may explain why these neurons have not been described previously, since it would be impossible to distinguish them from perivascular sympathetic neurons based upon their expression of Th. Importantly, we find no evidence for TrkC^CreERT2^ mediated recombination in sympathetic ganglia, while it is coexpressed with Th in DRG. This implies that TrkC^+^/Th^+^ nerves are indeed of sensory origin with their cell bodies in the spinal ganglia. This raises an important question as to whether Th has an enzymatic role in sensory neurons, and whether they can synthesize and release norepinephrine for a direct action on blood vessels. Our in vitro imaging data suggests that this is not the case, because both optogenetic and chemogenetic activation of the neurons did not elicit a change in blood vessel diameter in isolated tissue. Similarly, analysis of RNAseq data reveals that the other enzymes required for catecholamine synthesis are not detected in TrkC^+^/Th^+^ neurons (Extended Data Fig. 1k, Zeisel et al., 2018) consistent with previous reports (Brumovsky et al., 2006; Kummer et al., 1990; Price, 1985; Vega et al., 1991). Future studies, that utilize conditional knockout of Th in TrkC neurons for example, will determine what, if any, is the functional role of Th in these neurons.

Our in vivo live imaging data revealed robust effects of TrkC^+^/Th^+^ neuronal stimulation on blood flow and vessel diameter. Using LCSI and 3P imaging across spatial scales, we observed a substantial decrease in blood flow to the skin, and a reduction in vessel cross sectional area. This indicates that vasoconstriction provoked by activation of TrkC^+^/Th^+^ neurons may divert regional blood flow away from peripheral tissues. Accordingly, in LCSI experiments we observed an almost complete loss of the detectable vasculature upon stimulation that commenced in the distal toes and proceeded proximally over time. One consequence of reduced peripheral blood flow would be tissue hypoxia and acidosis leading to ischemic pain (Anitescu, 2018). Accordingly, we observed a profound mechanical hypersensitivity upon local stimulation of TrkC^+^/Th^+^ neurons that paralleled the reduction in vascularized area. Of note, alterations in neuronal control of vasculature have been associated with a number of painful conditions including diabetic neuropathy, migraine, Raynaud’s disease and fibromyalgia (Albrecht et al., 2013; Burnstock, 2008; Cooke and Marshall, 2005; Edvinsson et al., 2019; Jacobs and Dussor, 2016; Pietrobon and Moskowitz, 2013; Queme et al., 2017; Schmidt, 2014). To date, the majority of studies have considered only the contribution of autonomic and peptidergic sensory neurons to these conditions. However, the identification of a new class of non-peptidergic perivascular sensory neurons now opens up the possibility that they may also have a role in vascular associated pain. Identification of TrkC and Th as markers of these neurons will now enable further studies on their function, and help elucidate their role in regulating peripheral blood flow under basal conditions and in disease.

## Acknowledgments

We thank Philip Hublitz of EMBL Gene Expression Services, Pedro Moreira of EMBL Transgenic Services, and Violetta Paribeni for technical support of our work.

## Author contributions

CM, LC, SJB, LLS, AW, FJT, BC, AB, JS, TF, LS, BD and JS acquired data and performed analysis, KMB developed computational analysis methods for LSCI, RP, SGL, PAH gave feedback on experimental aspects, supervised experimental approaches, and implemented the data interpretation. LC prepared the figures. PAH designed the study with help from CM and LC and wrote the manuscript.

## Declaration of interests

The authors declare no competing interests.

## Methods

### EXPERIMENTAL MODEL AND SUBJECT DETAILS

#### Transgenic animals

##### Generation of TrkC^CreERT2^ mice

A bacterial artificial chromosome (BAC) containing the *TrkC* mouse locus was obtained from SourceBioscience (RP23-38E14) and a modified CreERT2-pA-Frt-Ampicillin-Frt cassette was inserted into the ATG of *TrkC*. The positive clones were confirmed by PCR and a full-length sequencing of the inserted cassette was performed. The ampicillin cassette was then removed using bacterial Flp and the accomplished removal was confirmed by sequencing analysis. Purified BAC DNA was then dissolved into endotoxin-free TE and prepared for intracytoplasmic sperm injection (ICSI). The method successfully produced offspring and the mice genotype was determined by performing PCR using the following primers: gcactgatttcgaccaggtt (fwd) and gagtcatccttagcgccgta (rev), yielding products of 408 bp.

##### Avil^hM3Dq^ mice

For gain of function studies, Avil^hM3Dq^ mice as described previously(Dhandapani et al., 2018a) were crossed to TrkC^CreERT2^ to generate TrkC^CreERT2^::Avil^hM3Dq^ heterozygous mice.

##### Rosa26^ChR2-YFP^ mice

For optogenetic experiments, TrkC^CreERT2^ mice were crossed to Rosa26^ChR2-YFP^ mice (The Jackson Laboratory, 024109) to generate TrkC^CreERT2^::Rosa26^ChR2-YFP^ mouse line.

##### Avil^iDTR^ mice

For diphtheria toxin-mediated ablation, Avil^iDTR^ mice, as described in (Stantcheva et al., 2016), were crossed to TrkC^CreERT2^ to generate TrkC^CreERT2^::Avil^iDTR^ heterozygous mice. Triple transgenic mice were also generated by crossing TrkC^CreERT2^::Avil^iDTR^ to Rosa26^ChR2-YFP^ mice. The obtained TrkC^CreERT2^::Avil^iDTR^::Rosa26^ChR2-YFP^ mice were used as a reporter line in ablation experiments. For all the experiments, littermates lacking Cre were used as controls.

All mice were housed in the EMBL Epigenetics and Neurobiology Unit, Rome, according to the Italian legislation (Art. 9, 27 Jan 1992, no 116) under licence from the Italian Ministry of Health and in compliance with the ARRIVE guidelines. Experimental protocols were approved by the EMBL Rome Ethics Committee and the Italian Ministry of Health.

### METHOD DETAILS

#### Tamoxifen treatment

To induce the expression of Cre, adult mice (older than 8 weeks of age) were treated intraperitoneally (i.p.) with 75 mg/kg of body weight of tamoxifen (Sigma Aldrich, T5648) dissolved in sunflower seed oil (Sigma Aldrich, S5007) for 3 consecutive days. Mice were then used for experiments at least one week after the last injection.

In some cases, to restrict the expression of Cre to DRG neurons, TrkC^CreERT2^::Rosa26^ChR2-YFP^ mice were treated with a single intrathecal (i.t.) injection of 45 ng of 4-hydroxytamoxifen (4-OH Tamoxifen, Sigma Aldrich, H7904). Experiments were performed at least one week after the treatment.

#### Electrophysiology

Whole cell patch clamp recordings from dissociated small TrkC^+^ DRG neurons were acquired with an EPC-10 double patch clamp amplifier (HEKA) controlled with Patchmaster© software (HEKA). Patch pipettes with a tip resistance of 2-5 MΩ were filled with intracellular solution 110 mM KCl, 10 mM NaCl, 1 mM MgCl2, 1 mM EGTA, 10 mM HEPES, 2 mM guanosine 5’-triphosphate (GTP) and 2 mM adenosine 5’-triphosphate (ATP) adjusted to pH 7.3 with KOH. The extracellular solution (ECS) contained 140 mM NaCl, 4 mM KCl, 2 mM CaCl2, 1 mM MgCl2, 4 mM glucose, 10 mM HEPES, adjusted to pH 7.4 with NaOH. Cells were clamped to a holding potential of −60 mV. Pipette and membrane capacitance were compensated using the auto function and series resistance was compensated to a 50%. For capsaicin and pH stimulation, clamped cells were perfused for 20s with ECS with 1 μM Capsaicin (Sigma) or ECS adjusted to pH 5.4. For mechanical stimulation, a series of mechanical stimuli in 1 μm increments with a fire-polished glass pipette (tip diameter 2-3μm) were applied at an angle of 45° and with a velocity of 1 μm/ms by a piezo driven micromanipulator (Nanomotor© MM3A, Kleindiek Nanotechnik). The evoked whole cell currents were recorded with a sampling frequency of 10 kHz and filter with 2.9 kHz low-pass filter. The mechanically activated currents were classified according to the inactivation kinetics obtained by fitting the current decay to a single exponential curve (C1+C2*exp(−(t−t0)/T) in rapidly adapting (RA, tau <10 ms) and intermediate adapting (IA, tau 15-30 ms). No slowly adapting currents (200-300 ms) were observed in the small TrkC^+^ population. For the classification of pH elicited currents, the inactivation kinetic was also calculated by fitting the current decay to a single exponential curve.

#### Immunofluorescence

Ganglia were dissected and fixed in 4%PFA overnight at 4°C. They were then embedded in 2% agarose (Sigma Aldrich, A9539) and cut in 50 μm sections using a vibratome (Leica, VT1000S). After an incubation of 30 minutes with a blocking solution containing 5% goat serum and 0.01% Tween-20 in PBS, the sections were incubated with one or more primary antibodies (Table 1) in blocking solution overnight at 4°C. As a negative control, sections were incubated without primary antibodies. The next morning, secondary antibodies in blocking solution were added and the sections were incubated for 1 hour and 30 minutes at room temperature (RT). After two washes with PBS, slides were mounted with prolong gold (Invitrogen, P36930).

**Table 1.** List of primary antibodies and lectins used for immunohistochemical experiments.

| Antibody | Concentration | Supplier/catalog number |
| --- | --- | --- |
| Rabbit anti-TH | 1:1000 | Millipore, AB152 |
| Mouse anti-CGRP | 1:500 | Rockland, 200-301-D15 |
| Rabbit anti-PV | 1:1000 | Swant, PV27 |
| Isolectin GS-B4-biotin XX-conjugate | 1:100 | Invitrogen, I21414 |
| Rabbit anti-desmin | 1:200 | Abcam, Ab32362 |

To examine TrkC^+^ peripheral afferents, mice were injected intravenously (i.v.) with a solution of 2% Evan’s Blue (Sigma Aldrich, E2129) in PBS. After 30 minutes, the skin was carefully dissected and fixed in 4% PFA overnight at 4°C. After permeabilizing with PBS-T (0.3% TritonX in PBS) for 30 minutes at RT, the tissue was incubated in a blocking solution (10% goat serum in PBS-T) for 2 hours at RT and then with one or more primary antibodies (Table 1) in blocking solution overnight at 4°C. Secondary antibodies were added in blocking solution for 1 hour and 30 minutes at RT and then the tissue was whole-mounted using prolong gold.

For immunofluorescence experiments, the following primary antibodies and lectins were used:

All secondary antibodies were Alexa-conjugated and were used at a concentration of 1:1000. Streptavidin-conjugated secondary antibodies were used at a concentration of 1:500.

All images were acquired using a Leica SP5 confocal microscope and analysed using ImageJ.

#### Ex-vivo live imaging

Mice were injected i.v. with a solution of 2% Evan’s Blue. After 30 minutes, the skin of the hind limb was carefully dissected and placed in a bath chamber where physiological conditions were maintained (32°C, 5% CO2, synthetic interstitial fluid: 108 mM NaCl, 3.5 mM KCl, 0.7 mM MgSO4, 26 mM NaHCO3, 1.7 mM NaH2PO4, 1.5 mM Cacl2, 9.5 mM sodium gluconate, 5.5 mM glucose and 7.5 mM sucrose at a pH of 7.4).

In the case of TrkC^CreERT2^::Avil^hM3Dq^ mice, 50 μM of Clozapine-N-oxide (CNO, Tocris, 4936) was added to the chamber. As a positive control, L-norepinephrine hydrochloride (Sigma Aldrich, 74480) was used at a concentration of 10 mM.

For TrkC^CreERT2^::Rosa26^ChR2-YFP^ mice, the skin was stimulated for 40 seconds every minute for 14 minutes with a built-in 488 nm microscope laser.

All tissues were imaged using a Nikon Ti Eclipse spinning disk confocal microscope. Images were acquired every minute for 14 minutes and analysed using ImageJ. For each blood vessel, the change in diameter was measured by randomly selecting three areas and comparing the initial diameter with the diameter at the end of the acquisition. The mean diameter change for each vessel was then expressed as percentage of the initial diameter.

#### AAV production and administration

Recombinant AAV-PHP.S (Chan et al., 2017) carrying Cre-dependent pAAV-hSyn-DIO-hM4D(Gi)-mCherry (Krashes et al., 2011) as a cargo (a gift from Bryan Roth (Addgene plasmid #44362; http://n2t.net/addgene:44362; RRID: Addgene_44362)) was produced in HEK293 cells as described previously (Grieger et al., 2006). Cells were harvested 5 days post infection, lysed with Triton X-100 at 0.5%, nuclease treated, concentrated by tangential flow filtration, and purified using isopycnic ultracentrifugation (Dias Florencio et al., 2015). Vector genome titration was performed using Q-PCR with primers targeting the promoter region of the viral cargo (Grieger et al., 2006). TrkC^CreERT2^ mice were injected at postnatal day 2 (p2) in the temporal vein using an Hamilton syringe with a 30G needle. 1e+12 viral genomes were injected per each pup. At 8 weeks of age mice were treated with tamoxifen as previously described and experiments were performed one week later.

#### Administration of DREADD ligands

In order to systemically activate TrkC^+^ neurons, TrkC^CreERT2^::Avil^hM3Dq^ mice were injected i.p. with 2.5 mg/kg of body weight of the DREADD agonist compound 21 dihydrochloride (C21, Hello Bio, HB6124).

For local activation or silencing, 2.5 mg/kg of CNO was injected subcutaneously in the hind paw of TrkC^CreERT2^::Avil^hM3Dq^ mice and TrkC^CreERT2^ injected with Cre-dependent AAV-hSyn-DIO-hM4D(Gi)-mCherry, respectively.

#### Propranolol administration

The nonselective β-adrenergic blocker propranolol hydrochloride (Sigma Aldrich, P0884) was injected i.p. at a concentration of 5 mg/kg of body weight. The injection was performed immediately after the administration of the DREADD ligand.

#### Diphtheria toxin injection

TrkC^CreERT2^::Avil^iDTR^ mice were injected i.p. with 40 ng/g of body weight of diphtheria toxin (DTX, Sigma Aldrich, D0564). All mice were monitored during the injection period and blood pressure, heart rate and blood flow were measured before the injection of DTX and 16, 24 and 32 hours after.

#### Blood pressure measurements

Mice were anesthetized with a 2% isoflurane and medical air mixture through a nose cone and placed on a heat pad at 37°C. Blood pressure (BP) was measured using a Non-Invasive Blood Pressure (NIBP) system (AD Instruments) paired with a PowerLab 4/20 ML840 (AD Instruments) and LabChart 4 software to acquire and analyze data. For each measurement, BP was registered four times per mouse with a 1-minute interval and the mean value was recorded.

#### Heart rate measurements

Mice were anesthetized with 2% isoflurane and kept at 37°C using a heat pad. The heart rate was monitored for 30 or 60 minutes using a PhysioSuite MouseSTAT (Kent Scientific) and Free Serial Port Terminal 1.0.0.710 software. All data were analysed using gHRV 1.6 software(Rodriguez-Linares et al., 2014).

#### Laser Speckle Contrast Imaging

To analyse blood flow, recordings were performed using a 780 nm, 100mW laser (LaserLands) at a working distance of 5 cm and a Leica Z16 Apo microscope with a high resolution camera (AxioCam MRM, Carl Zeiss) with 5 ms exposure time at maximum speed for 100 cycles. Data were then analyzed using a custom Matlab script (Gnyawali et al., 2017), and vascularized area analysed using ImageJ (mm^2^ occupied by blood vessels in the field of view at each time point).

For gain of function experiments, mice were anesthetized with an i.p. injection of 90 mg/kg ketamine (Lobotor, Acme) and 0.5 mg/kg medetomidine (Domitor, Orion Pharma) and their hind paw was attached using double-sided tape to a plastic platform. Images were acquired before the treatment with CNO and after its administration every 2 minutes for 30 minutes.

For ablation experiments, mice were anesthetized with 2% isoflurane and their ear was attached to a plastic platform with the external side up, facing the camera. Images were acquired before the injection of DTX and 16, 24 and 32 hours after.

#### In-vivo three-photon (3P) imaging

To analyze blood flow and blood vessel cross-section changes *in-vivo*, we performed 3P microscopy using a custom-built system. 3P-excitation of fluorescein labeled blood plasma was done at 1300nm by using a non-collinear optical parametric amplifier (NOPA, Spectra Physics) pumped by a regenerative amplifier (Spirit, Spectra Physics) with a repetition rate of 400kHz. Dispersion compensation to maintain short pulse durations (50 fs) at the sample plane was achieved with a custom-built dispersion compensation unit (Lingjie Kong and Cui, 2013). For imaging the power under the objective lens (Olympus 25x, 1.05NA, XLPLN25XWMP2) ranged from 8mW to 12mW.

To label blood vessels *in-vivo*, mice were injected with 150μl of 5% w/v fluorescein-dextran (Sigma FD2000S 2MDa). For experiments mice were anesthetized with 2% isoflurane and the hair of the ear was removed using hair removal crème. The ear was attached to a cover glass using double sided tape with the external side facing the objective lens.

For optogenetic stimulation, an area of the ear of TrkC^CreERT2^::Rosa26^ChR2-YFP^ or control mice with a diameter of 2mm was illuminated with a 470nm LED (Thorlabs M470L4) for 1min. Stimulation consisted of 10ms bursts at 5Hz with 3mW/mm^2^ excitation power.

For volume/stack data analysis, first a median filter (3×3 kernal; ImageJ) was applied before manual segmentation of the blood vessel. A maximum intensity image was generated from the segmented stack and the cross-sectional area of the blood vessel was measured using the ‘analyze particles’ plugin from ImageJ. For visualization purposes of the orthogonal views the image stack was interpolated in the axial dimensions by a factor 5.

In general, 3P imaging data was acquired at 0.69Hz, and typically 20 frames were averaged to improve signal-to-noise, manually segmented and the area analyzed using the ‘analyze particles’ plugin from ImageJ.

#### Behavioural testing

All behaviour experiments were performed on adult male mice (>8 weeks of age). Littermates not expressing Cre were used as controls. The experimenter was always blind to the genotype of the mice. Unless otherwise specified, all tests were performed 1 hour after local injection of CNO.

##### Acetone drop test

Mice were habituated on an elevated platform with a mesh floor for 30 minutes. A single drop of acetone was sprayed on the hind paw with a blunt syringe making sure not to touch the paw. The test was repeated 5 times per mouse and the behavioural responses were scored as follows:

0 = no response
1 = paw withdrawal or single flick
2 = repeated flicking
3 = licking of the paw

##### Paintbrush test

After a habituation time of 30 minutes on an elevated platform with a mesh floor, the hind paw of mice was stimulated with a paintbrush in the heel-to-toe direction. The responses were scored according to Duan et al. 2014 (Duan et al., 2014):

0 = no response
1 = paw withdrawal
2 = flicking of the paw
3 = licking of the paw

##### Von Frey test

Mice were placed on an elevated platform with a mesh floor and habituated for 30 minutes. The hind paw was stimulated with calibrated von Frey filaments (North coast medical, NC12775-99) and the 50% withdrawal thresholds were calculated using the Up-Down method previously described (Chaplan et al., 1994).

To measure the sensitivity to mechanical pain over time, the hind paw of TrkC^CreERT2^::Avil^hM3Dq^ mice was stimulated with a 0.02 g filament five times, 10 minutes, 25 minutes and 40 minutes after the local injection of CNO. The percentage of withdrawals was calculated per each time point.

##### Hot plate test

To measure thermal nociception over time, mice were placed on top of a hot plate (Ugo Basile, 35150) set at 52°C and the latency to response, indicated as flicking or licking of the hind paw, was measured before local injection of CNO in the hind paw and 10, 25 and 40 minutes after.

### QUANTIFICATION AND STATISTICAL ANALYSIS

All statistical data are represented as standard error of the mean (SEM). Student’s t-test and/or 2-way repeated measures ANOVA were used and p<0.05 was considered statistically significant.

## Supplemental Information

**This file includes:**

Supplementary Figures S1 to S4

Captions for Videos S1 to S7

**Other Supplementary Materials for this manuscript include the following:**

Videos S1 to S7

**Figure S1.**
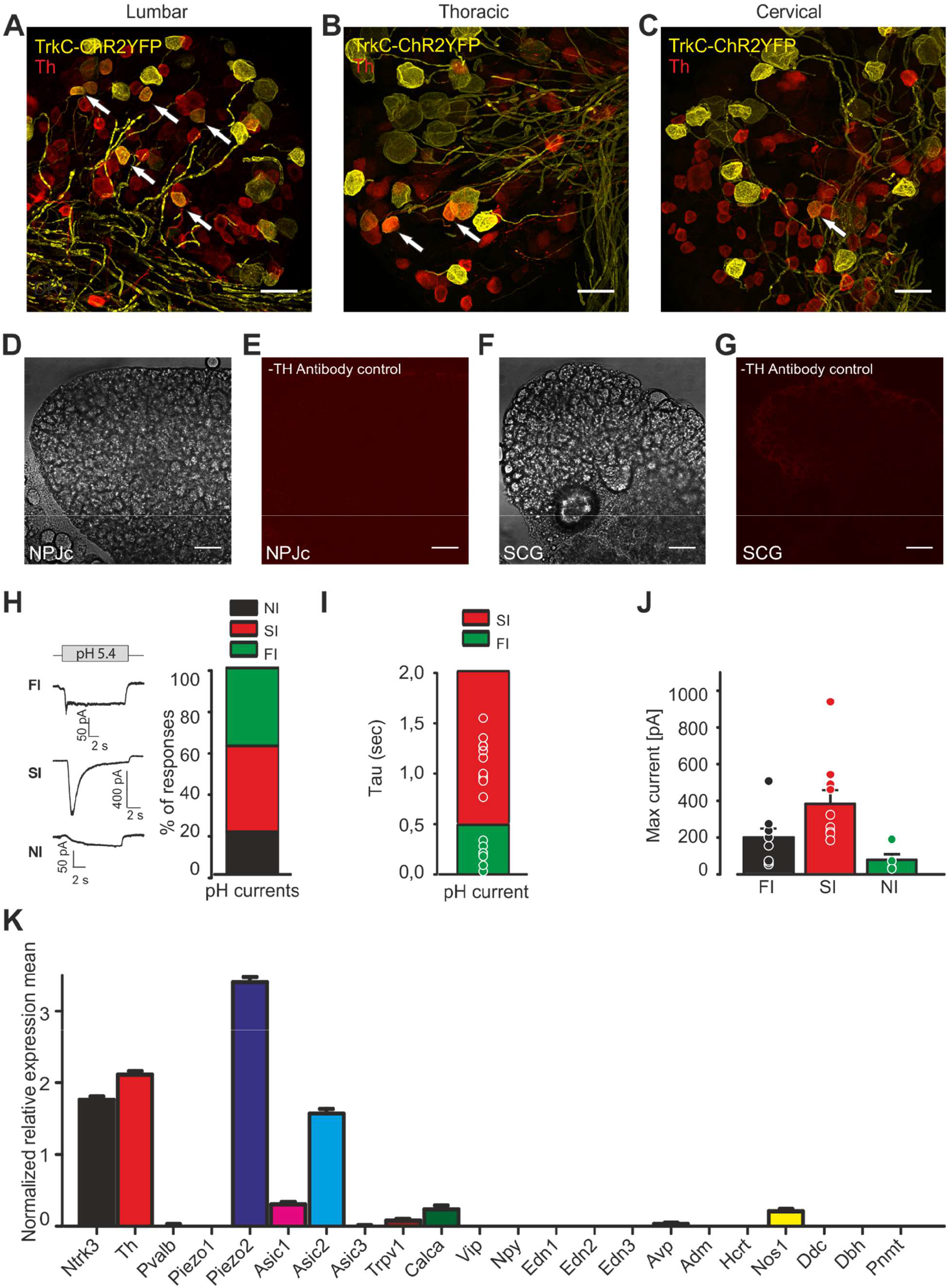
Characterization of TrkC^+^ neurons. (A-C) Immunofluorescent staining of DRG sections from TrkC^CreERT2^::Rosa26^ChR2-YFP^ mice labelled with anti-Th antibodies. TrkC expression was investigated in lumbar DRG (A), thoracic DRG (B) and cervical DRG (C). Double positive neurons are indicated by arrows. Scale bars, 50 μm. (D-G) Antibody controls for TH showing phase contrast (D and F) and red channel images(E and G) in the NPJc and SCG. (H) *Left*, example traces of pH activated currents recorded in small TrkC^+^ neurons (24 cells from 3 mice). *Right*, distribution of neurons according to the pH current type: fast inactivating (FI), slowly inactivating (SI) and non-inactivating (NI). (I) Distribution of the inactivation constants of the pH elicited currents. (J) Comparison of the maximal current amplitude in the different pH elicited current types. (K) Gene expression panel of TrkC^+^ Th^+^positive neurons (n=197).

**Figure S2.**
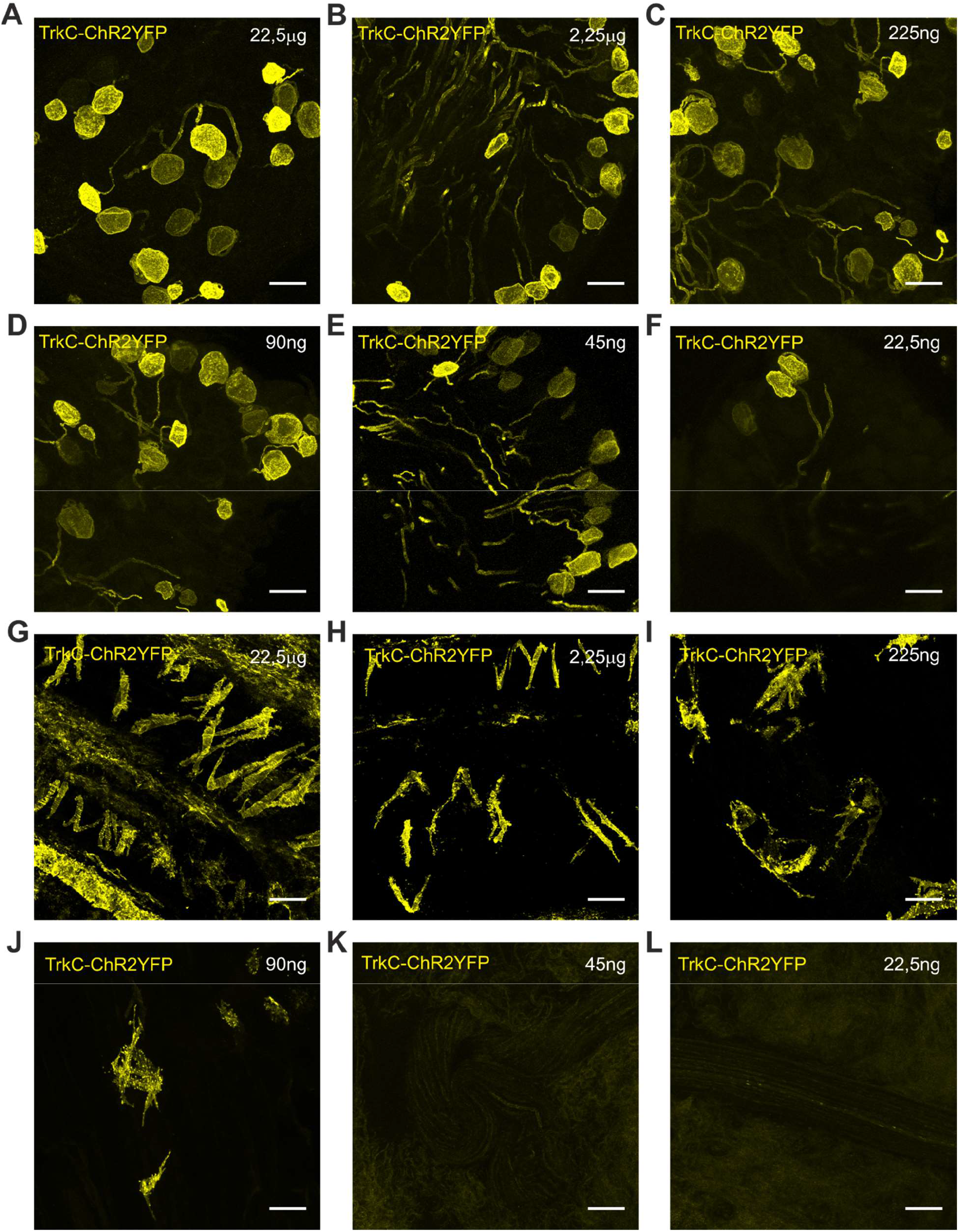
Concentration dependent recombination upon intrathecal injection of 4-OH tamoxifen. (A-F) Cre-driven recombination and YFP expression in DRG neurons and vSMCs. (G-L) At lower concentrations, YFP is evident only in DRG and not in vSMCs. Scale bars 50 μm.

**Figure S3.**
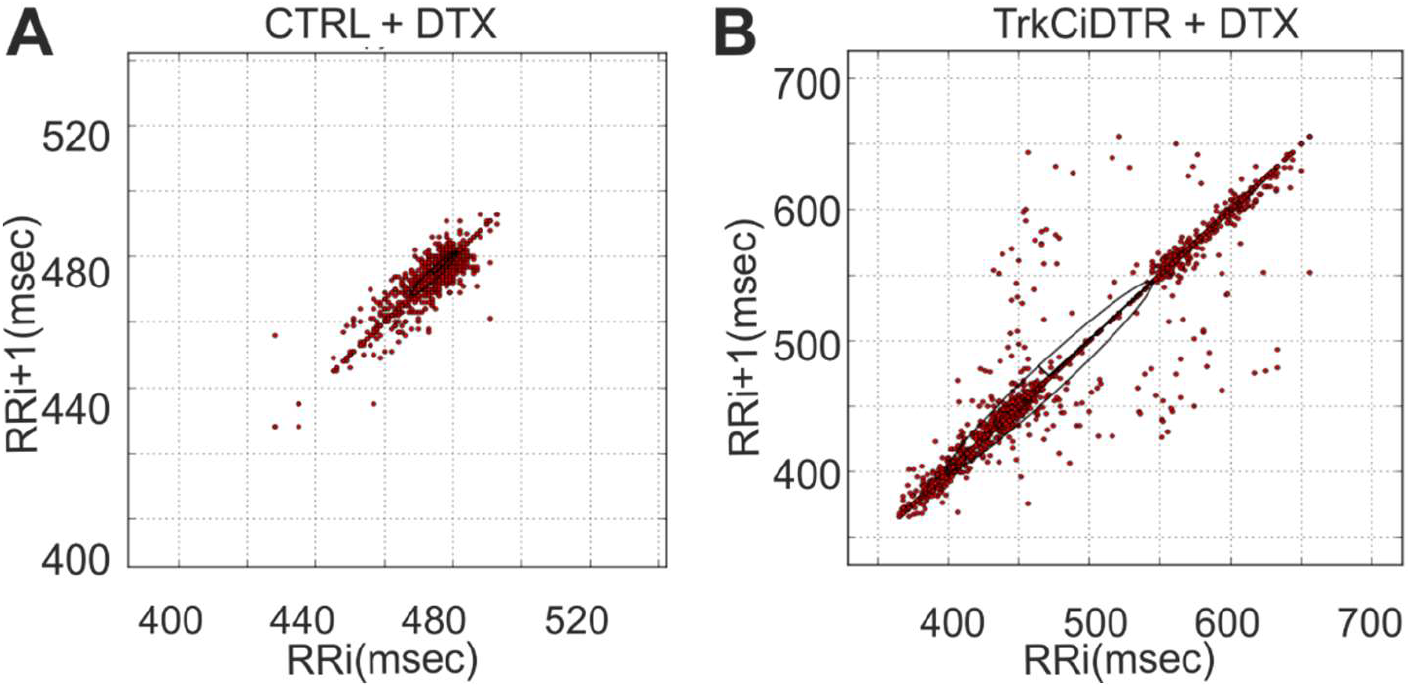
Ablation of TrkC^+^ neurons. (A and B) Representative Poincaré plots of a control (A) and TrkC^CreERT2^::Avil^iDTR^ (B) mouse treated with DTX.

**Figure S4.**
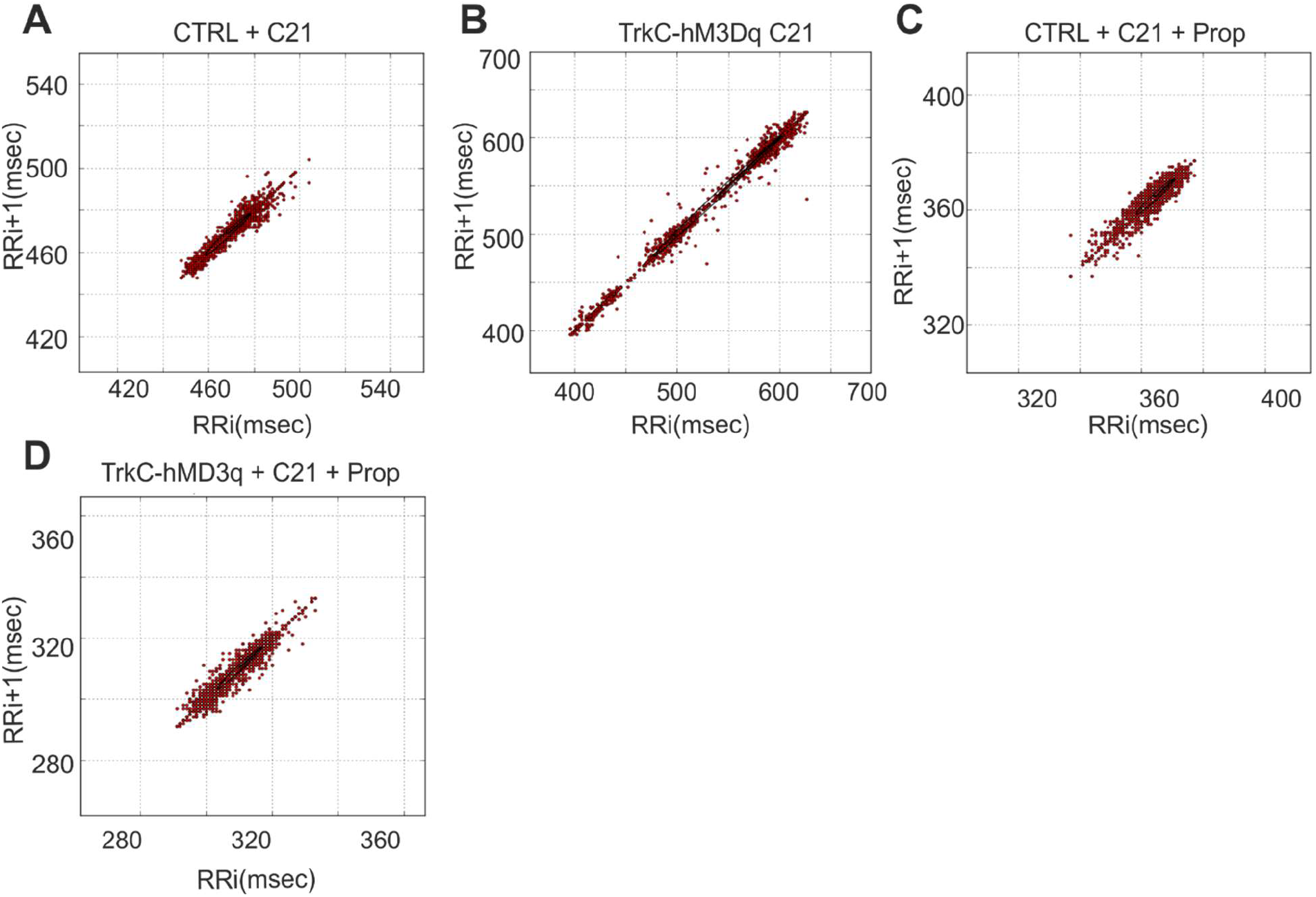
Activation of TrkC^+^ neurons. (A and B) Representative Poincaré plot of a control (A) and a TrkC^CreERT2^::Avil^hM3Dq^ (B) mouse treated with C21. (C and D) Representative Poincaré plot of a control mouse treated with C21 and Propranolol (C) and a TrkC^CreERT2^::Avil^hM3Dq^ mouse treated with C21 and Propranolol (D).

### Supplementary Video Captions

**Video S1.** 3-photon image time series of fluorescein labelled blood plasma in the ear of a TrkC^CreERT2^::Rosa26^ChR2-YFP^ mouse post-stimulation. (Frame rate 0.67Hz, Field-of-View 500μm). A decrease in vessel cross-sectional area can be observed.

**Video S2.** 3-photon image time series of fluorescein labelled blood plasma in the ear of a TrkC^CreERT2^::Rosa26^ChR2-YFP^ mouse pre-stimulation. (Frame rate 0.67Hz, Field-of-View 500μm)

**Video S3.** 3-photon image time series of fluorescein labelled blood plasma in the ear of a TrkC^CreERT2^::Rosa26^ChR2-YFP^ mouse post-stimulation. A reduction in blood flow can be observed. (Frame rate 0.67Hz, Field-of-View 500μm).

**Video S4.** Chemogenetic stimulation of TrkC^CreERT2^::Avil^hM3Dq^ neurons with CNO does not lead to any change in blood vessel diameter.

**Video S5.** Optical stimulation of TrkC^CreERT2^::Rosa26^ChR2-YFP^ neurons does not provoke any change in blood vessel diameter.

**Video S6.** Optical stimulation of TrkC^CreERT2^::Rosa26^ChR2-YFP^ vSMCs provokes blood vessel shrinkage.

**Video S7.** Application of norepinephrine caused a statistically significant reduction in vessel diameter.

